# Lipid distributions and transleaflet cholesterol migration near heterogeneous surfaces in asymmetric bilayers

**DOI:** 10.1101/2021.01.04.425270

**Authors:** Elio A. Cino, Mariia Borbuliak, Shangnong Hu, D. Peter Tieleman

## Abstract

Specific and nonspecific protein-lipid interactions in cell membranes have important roles in an abundance of biological functions. We have used coarse-grained (CG) molecular dynamics (MD) simulations to assess lipid distributions and cholesterol flipping dynamics around surfaces in a model asymmetric plasma membrane containing one of six structurally distinct entities: aquaporin-1 (AQP1), the bacterial β-barrel outer membrane proteins OmpF and OmpX, KcsA potassium channel, WALP23 peptide, and a carbon nanotube (CNT). Our findings revealed varied lipid partitioning and cholesterol flipping times around the different solutes, and putative cholesterol binding sites in AQP1 and KcsA. The results suggest that protein-lipid interactions can be highly variable, and that surface-dependant lipid profiles are effectively manifested in CG simulations with the Martini force field.

## Introduction

Biological membranes are fundamental to life as we know it. They regulate signaling, compartmentalization, and the flow of ions and other substances. Proteins are an integral component of cellular membranes, and have essential roles in conferring distinct functionalities, which are often mediated through interactions with lipid molecules.^1,2^ Prominent examples of biochemical process that involve protein-lipid interactions include regulation of sperm-egg fusion by lipid anchoring of CD9/CD81,^3^ and maintenance of lung homeostasis via association of surfactant proteins with cholesterol and phospholipids.^4^

Although protein-lipid interactions are known to be involved in numerous biological processes, their experimental characterization can be a challenge, and MD simulations at all-atom (AA) and CG approximations have become important tools for their characterization.^5^ While AA models can resolve molecular details more precisely, CG models can provide improved sampling of rather slow processes, such as equilibration of lipid distributions, and cholesterol movement between leaflets.^6^ The lipid profiles in the vicinities of integral membrane proteins can be important for their functionality, and MD simulations have proven useful in studying the association between proteins and specific lipid types.^7–9^ Membrane lipids often organize into distinct microdomains, or lipid rafts, of varying compositions around different proteins.^10^ Using CG simulations, it has been shown that the phospholipid and cholesterol distributions in a membrane can be specific to GPCR type and conformational state.^11^ In addition to lateral movement within monolayers, lipids can also transition between leaflets, which can be induced by the presence of certain proteins.^12^ For example, ABCA1 specifically promotes cholesterol translocation from the inner to outer leaflet for regulation of cellular events without altering the total cholesterol content in the membrane.^13^ Flips between leaflets occurs relatively slowly for phospholipids, but sterols, such as cholesterol, can exchange between leaflets on the microsecond timescale, which can be studied using MD simulations at the CG level.^14^

Here, we performed a systematic comparison lipid distributions and cholesterol flipping in an asymmetric bilayer around six distinct surfaces, including eukaryotic and bacterial proteins, as well as synthetic entities (Fig. 1). Though most bacteria do not synthesize cholesterol, some can obtain it from their environment, and many can synthesize sterol-like lipids, which may act as functional analogs.^15^ Nevertheless, the different entities included in the study were primarily selected to encompass a range of chemical and structural diversity to identify features of lipid distributions that have a generic basis versus a possible functional importance. The systems were designed to be potentially useful in the future as model systems to test the agreement (or lack thereof) between Martini simulations, the new Martini 3 version, further development of Martini 3, and atomistic simulations. Variations in lipid localization and cholesterol flipping dynamics were observed around the different surfaces in the individual leaflets, suggesting that protein-lipid interactions can be highly variable, and that unique lipid environments may be recruited by specific proteins.

**Fig. 1.**
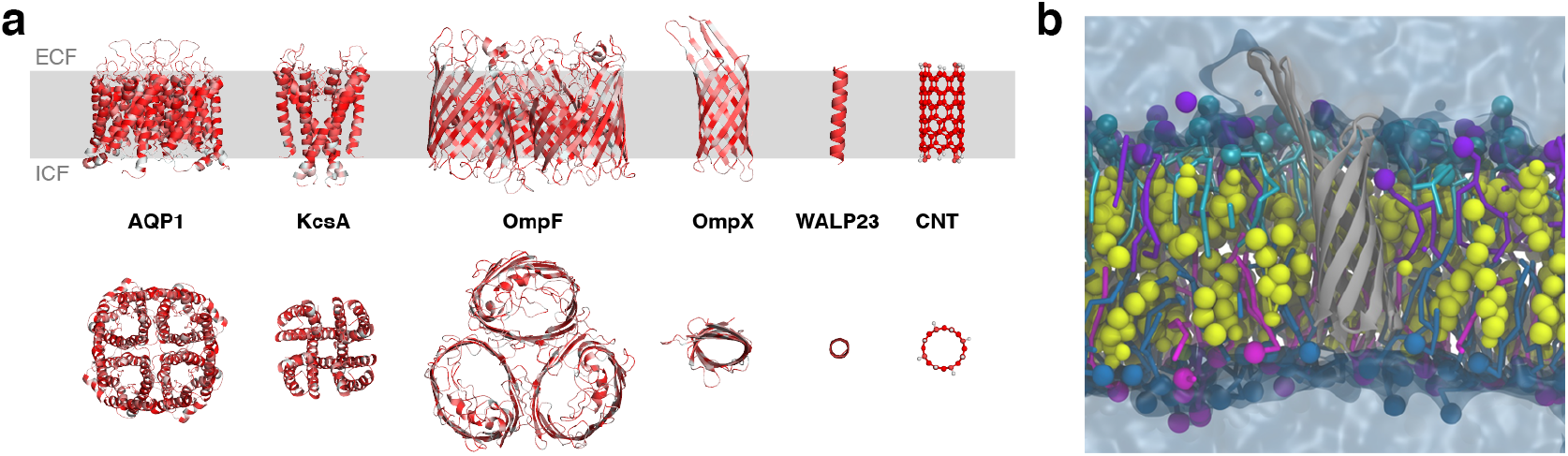
Asymmetric model bilayer embedded with different membrane proteins and a carbon nanotube. **a** Membrane proteins and carbon nanotube models. AQP1, KcsA, OmpF, OmpX, and WALP23 peptide are shown in cartoon style, and CNT with ball and stick, colored from low to high hydrophobicity on a white-red gradient. The upper and lower panels show side and a top view, respectively. ECF: extracellular fluid; ICF: intracellular fluid. **b** Initial state of the OmpX system: OmpX (gray), CHOL (yellow), POPC (purple), DPSM (cyan), POPE (blue), POPS (magenta), and water (translucent).

## Methods

### System construction

Solvated asymmetric bilayer systems and topologies were built using the Martini Maker^16^ module of CHARMM-GUI^17^ and INSANE (INSert membrANE),^18^ with the Martini v2.2^19^ parameter set and nonpolarizable water. The systems were composed of 1-palmitoyl-2-oleoyl-phosphatidylcholine (POPC) and N-stearoyl-D-erythro-sphingomyelin (DPSM) at 1:1 ratios in the upper leaflet, and 1-palmitoyl-2-oleoyl-phosphatidylethanolamine (POPE) and 1-palmitoyl-2-oleoyl-phosphatidylserine (POPS) at ~2:1 ratios in the lower leaflet, with equal numbers of cholesterol molecules in both leaflets, and one of AQP1-1j4n,^20^ KcsA-1r3j,^21^ OmpF-3pox,^22^ OmpX-1qj9,^23^ WALP23 peptide, or CNT (Table S1). Two lipids only systems, one with half the number of cholesterol molecules, were also built (Table S1). The lipid composition used approximates that of a simplified mammalian red blood cell membrane.^24^ To conserve protein structure, an elastic network was applied on atom pairs within 0.9 nm with a 500 kJ mol^−1^ force constant.^25^ A Martini-compatible CNT structure and topology was generated using cnt-martini.^26,27^ The CNT consisted of 12 rings, 8 beads per ring, giving a length of 4.5 nm and diameter of 1.2 nm. For the top and bottom rings, the CNP beads were substituted for more polar SNda beads. Atomistic AQP1 systems and topologies were generated by backmapping the final frame of each replica using CHARMM-GUI, CHARMM36m force field,^28^ and TIP3P water model (Table S1).

### MD simulations

Energy minimization, stepwise equilibration, and final production runs at 310 K were carried out with GROMACS 2018^29^ using the default CHARMM-GUI generated run parameter files. Integration time steps of 20 and 2 fs were used for the CG and AA simulations, respectively. Each CG system was simulated in triplicate for 60 μs (3 × 20 μs), while 3 × 1 μs runs were conducted for the AA AQP1 systems.

### Data analysis

Considering the periodicity of the simulation box, all systems were centered around the protein or CNT center of mass, and aligned to remove rotation and translation. The gmx select tool was used to dynamically compute the number of lipids around the different solutes using the PO4 beads for POPC, DPSM, POPE, and POPS, and ROH beads for CHOL. Cholesterol flipping between the upper and lower leaflets was assessed by calculating the angle between the C2-ROH vector of the cholesterol molecule and the bilayer normal. Successful flips were those that exhibited a stable vector sign change and angle difference >150° for at least 10 ns. Average flipping time for each cholesterol was measured as the time duration between flipping events, and plotted as a function of distance to protein or CNT. Unless shown otherwise, data are presented as an average of the three replicas for each system over the second half (last 10 μs) of the trajectories. VMD was used for system visualization.^30^ Numpy and Matplotlib were used for statistical analysis and data plotting.^31,32^

## Results

To assess system equilibration and reproducibility across replicas, the number of contacts between membrane lipids and embedded proteins or CNT were calculated throughout the simulations (Fig. S1 and S2). Lipid distributions around the different surfaces tended to stabilize early on in the simulations, within a few microseconds, and there was good consistency between replicates. Although the cholesterol molecules in each system were evenly distributed between monolayers in the initial configurations, within a few microseconds of simulation, cholesterol became enriched in the upper leaflet, remaining stable thereafter around a 1.3:1 ratio, and 1.6:1 for the lipids only system with a halved cholesterol concentration (Table S2).

Next, the localization probability density of CHOL, POPC, DPSM, POPE, and POPS was computed for each system (Fig. 2a). Enrichment of cholesterol was observed around the surface of AQP1 in both leaflets, deeper within the KcsA structure in the upper leaf, and around the OmpF surface in the upper leaflet (Fig. 2a and b). There was a higher localization probability of POPC around AQP1, and at the trimer interfaces of OmpF, which was not evident in the other systems. POPS was enriched around the outer surfaces of AQP1 and KcsA in the lower leaflet, and increased POPE density around OmpF in the lower leaflet was noted (Fig. 2a). Aside from the aforementioned cases, lipid concentrations around the different surfaces tended to approximate their bulk values. The equilibrated lipid distributions around AQP1 were maintained in the atomistic continuation simulations (Fig. S3).

**Fig. 2.**
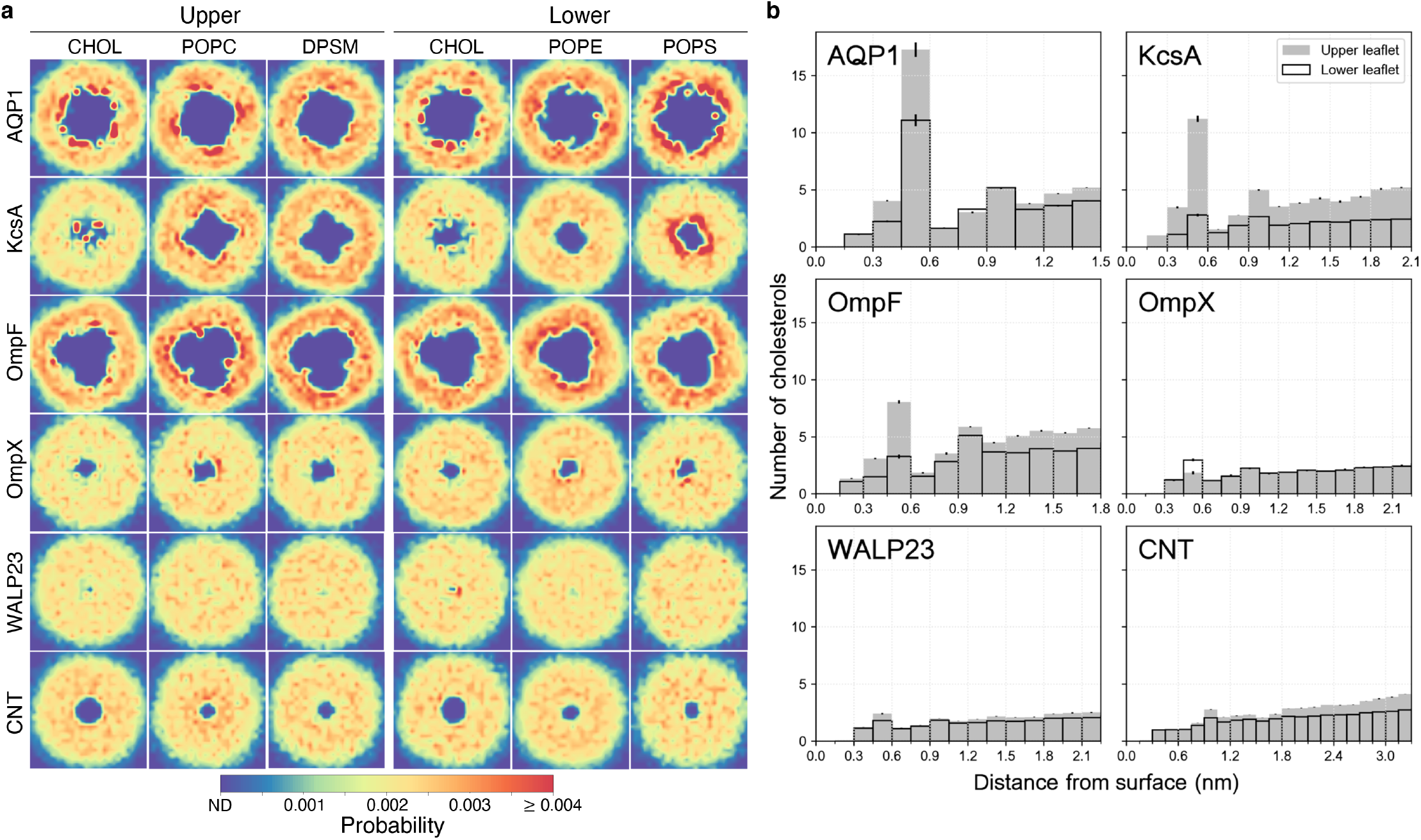
Lipid distributions. **a** Probability density of CHOL, POPC, DPSM, POPE, and POPS in the upper and lower leaflets. The areas in blue in the center of 2D maps show membrane regions displaced by proteins or CNT. **b** Cholesterol counts as a function of distance from different surfaces. The number of cholesterol molecules were counted as a function of nearest distance from the protein or CNT beads to cholesterol ROH beads for upper (shaded bar charts) and lower (empty bar charts) leaflets separately. Standard deviation is shown with black error bars.

Subsequently, per-molecule and overall cholesterol flipping was assessed. The different systems presented similar bulk flip times of ~1 μs, but slower flipping dynamics were evident close to AQP1 and KcsA (Fig. 3a and b). Several cholesterol molecules around AQP1 and KcsA had flip times >10 μs (Fig. 3a), indicating that they did not flip during the simulation interval analyzed. In the lipids only system with half the number of cholesterol molecules, flipping was faster, with an average rate of 1.4 flips/μs (Table S2). A few cholesterol molecules with flip times >1 μs were observed in the other systems at varying distances to the different surfaces, and also in the lipids only configurations, and was attributed to stochastic behavior (Fig. 3a and S4). No cholesterol flipping events were registered in the AQP1 AA systems.

**Fig. 3.**
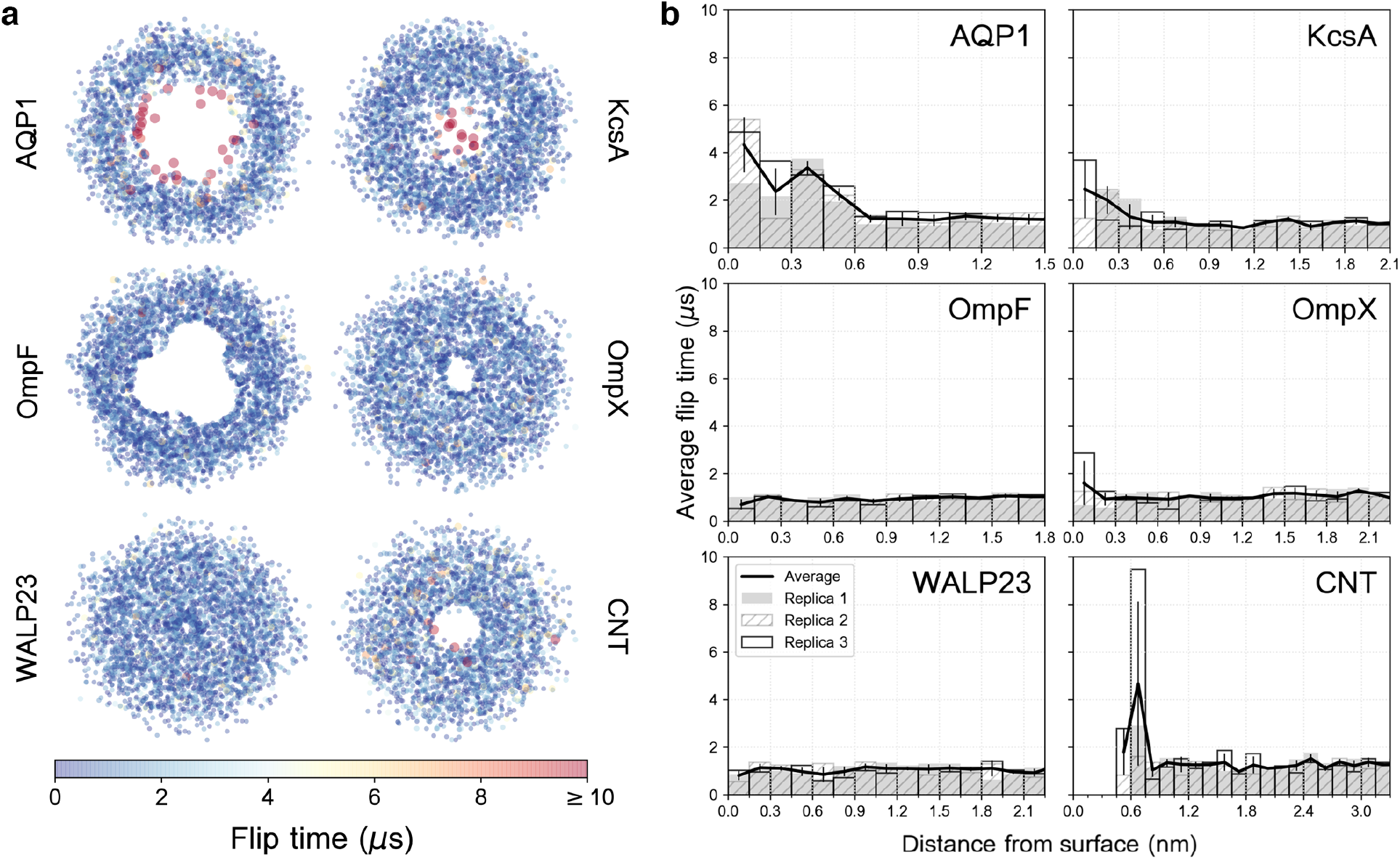
Cholesterol flip times around different surfaces. **a** Each spot represents a cholesterol flip event, with point size and color indicating the time from the previous flip. Each plot is centered around the protein’s center of mass. **b** Average cholesterol flipping times as a function of distance from the surfaces.

Because areas of higher CHOL density colocalized with those of slow flipping in AQP1 and KcsA, these regions were inspected more closely to assess possible cholesterol interaction sites. AQP1 and KcsA are similar in that they are both predominantly α-helical homotetramers. Analyzing the non-flipping cholesterol molecules from the AQP1 and KcsA trajectories, it was evident that they preferentially bind at the subunit interfaces (Fig. 4). Each intersubunit interface typically accommodated a single cholesterol molecule; however, localization to intersubunit groves was reproducible across different replicas, leading to the overlap of molecules in Fig. 4.

**Fig. 4.**
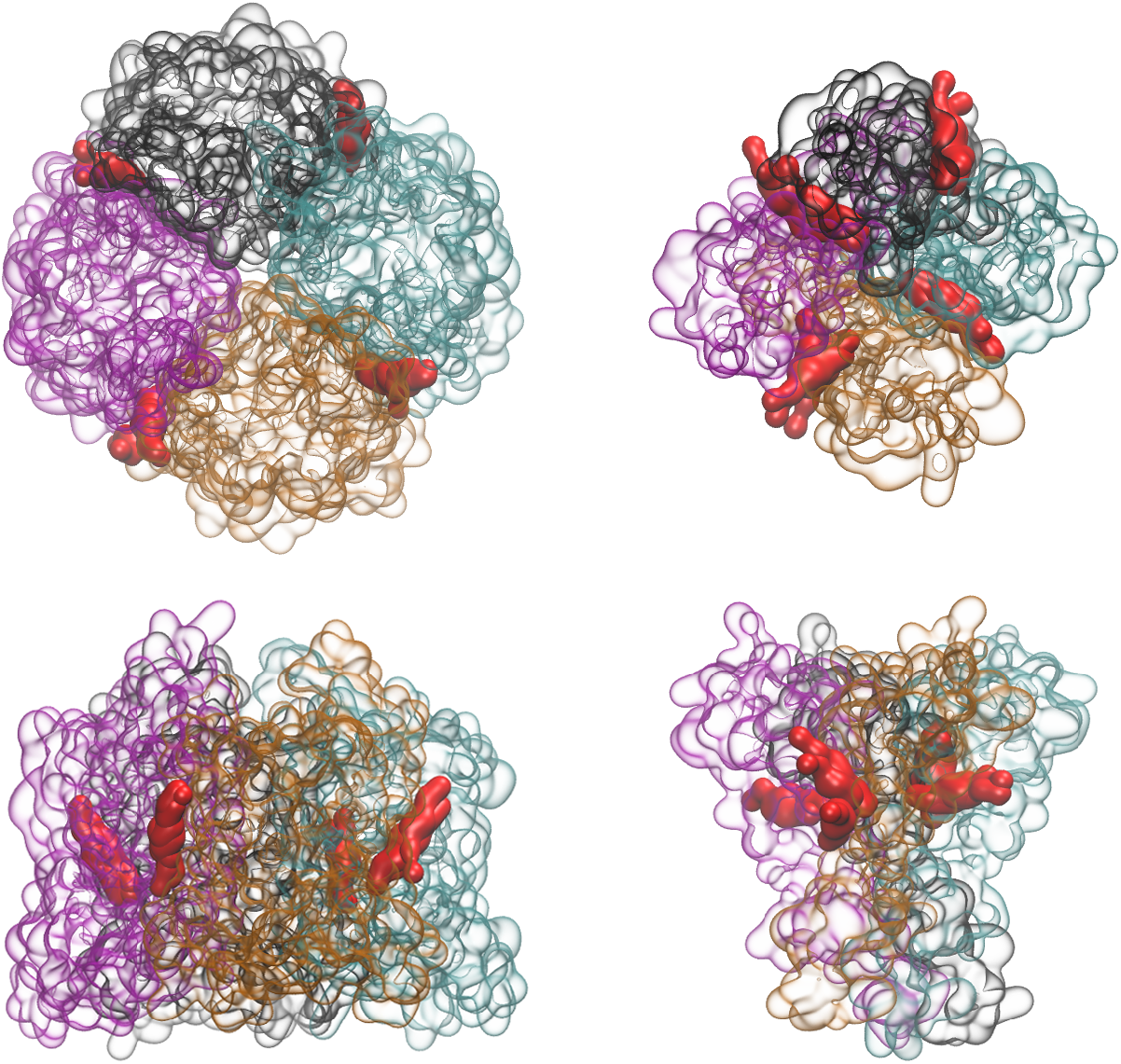
Locations of non-flipping cholesterol molecules around AQP1 and KcsA. Top and side views of non-flipping cholesterol molecules (red) from the AQP1 (left) and KcsA (right) trajectories. Individual subunits are colored in black, cyan, orange, and magenta.

## Discussion

Achieving a better understanding of lipid distributions and cholesterol dynamics around membrane proteins is crucial given the importance of protein-lipid interplay in biological processes.^33^ We have taken a methodical approach to compare per-leaflet lipid distributions and transleaflet cholesterol flipping around six different surfaces in an asymmetric model membrane. The results are accordant with previous reports of unique lipid environments around certain proteins. Here, follows a contextualized discussion of our observations.

Though the simulations were initiated with an equal number of cholesterol molecules in the upper and lower leaflets, reproducible migration of cholesterol from the lower to upper leaflet occurred in the early stages of the different simulations (Fig. S1 and S2 and Table S2). Similar behavior has been reported for asymmetric bilayers of comparable makeup, and possible explanations include cholesterol preference for certain lipid types, and leaflet coupling to equilibrate membrane stress.^34^ Allender and colleagues noted a similar degree of cholesterol enrichment in the outer leaflet as observed in our simulations, which they attribute to a strong attraction between cholesterol and sphingomyelin.^35^ They also predict that nearly 80% of the cholesterol could localize in the outer leaf if it were not for unfavorable bending energies associated with the spontaneous curvature induced by such a distribution. Our results showed that the saturated sphingomyelin lipid environment of the upper leaflet contained 25-40% more of the total cholesterol than the lower leaflet, with greater differences close to protein surfaces in some cases (Table S2 and Fig. 2). In the systems with the same lipid composition, but half the number of cholesterol molecules, the upper leaflet contained an average of 60% more cholesterol than the lower leaflet (Table S2). Based on the work of Allender et al., it seems reasonable that by reducing the total amount of cholesterol in the bilayer, the upper leaflet is able to accommodate a higher fraction of cholesterol without inducing excessive curvature. The ability of cholesterol to exchange between monolayers has been shown to eliminate membrane stress when it solvates equally well in both leaflets, and create stress, resulting in membrane curvature, when partitioning to one leaflet is preferred.^36^ It is evident that the intrinsic asymmetry of biomembranes, coupled with the ability of membrane components, such as cholesterol to undergo transleaflet migration leads to a complex, but biologically relevant situation. For example, it has been suggested that Caveolin-1 can induce membrane curvature through cholesterol recruitment, and that such caveolar complexes have central roles in cell signalling.^37^

As reported in other works,^1,11^ enrichment of specific lipid types around certain surfaces was detected in our simulations. One of the most evident observations was a high probability density of cholesterol around AQP1 and KcsA (Fig. 2), in particular at subunit interfaces (Fig. 4). Mediation of protein function through specific interactions with cholesterol and other lipids has been shown for several proteins, such as Kir channels and GPCRs.^38^ Specific interactions between cholesterol and aquaporins has been speculated to regulate their water permeabilities, though the mechanism by which this occurs has not been clarified.^39^ For KcsA, it has been shown that cholesterol and other sterols can attenuate ion conductance by regulating bilayer tension.^40^ In line with our findings, docking experiments indicate preferential binding of cholesterol to subunit interfaces on KcsA.^41^ It was suggested that cholesterol may compete with anionic lipids for KcsA binding sites, and this idea is supported by our finding of POPS enrichment around KcsA in the lower leaflet. Similarly, the activity of aquaporins can be regulated by the density of negatively charged lipids in their immediate vicinity,^42^ and we detected POPS clustering at the AQP1 surface in the inner leaflet (Fig. 2a). Additionally, POPC was found to associate at distinct sites around AQP1 in the upper leaflet, but further studies are required to assess the biological relevance of these interactions. A distinct lipid profile around OmpF was also noted (Fig. 2a), and our findings are consistent with experimental demonstration of POPE and POPC binding to OmpF and regulating its function.^43,44^ The OmpX, WALP23, and CNT systems presented considerably more homogeneous lipid distributions, with no clearly discernable patterns of enrichment around their surfaces (Fig. 2a). The absence of a particular lipid profile around the CNT was to be expected considering that its surface was not functionalized such that especially strong interactions would occur with other system components. However, in future studies it could be useful to evaluate if CNTs with different functional groups on their surface could recruit specific lipids, as such systems could serve as models for understanding the molecular details of lipid raft assembly. WALP peptides were developed as models for studying protein insertion in membranes, orientation, and hydrophobic mismatch.^45^ Although WALP23 did not present a distinct lipid profile in its vicinity, association with other lipid types has been demonstrated, suggesting that WALP and related KALP peptides could be useful models for studying protein-lipid interactions.^46^

Another property analyzed from our simulations was cholesterol flipping dynamics between leaflets. As discussed earlier, transleaflet migration of cholesterol can be influenced in a protein-dependant manner, regulating bilayer properties and biological functions. Computer simulations have shown that the rate of cholesterol flip-flop depends on several factors, such as bilayer composition, temperature, and force field.^14,47^ Because our aim was to compare systems of the same lipid composition, the absolute rates of interleaflet diffusion are not highly important, though flipping was found to occur on the timescale of a few microseconds, which is in line with other Martini simulations.^48^ It is also noteworthy that reducing the amount of cholesterol by half led to ~1.5x faster flipping (Table S2). Like the lipid distribution profiles, the most obvious differences in cholesterol flip time was seen in the vicinity of AQP1 and KcsA (Fig. 3). Slower transleaflet movement of cholesterol around these proteins was attributed to its tendency to associate at putative cholesterol binding sites at subunit interfaces (Fig. 4).

Finally, we have shown that CG simulations can be beneficial for equilibration of lipid profiles for subsequent atomistic studies. Due to the increased computational time needed to simulate atomistic versus coarse-grained systems, it can be efficient to perform initial system equilibration at the CG level. To illustrate this point, the last frames of the AQP1 systems simulated at the CG level were backmapped to atomistic detail, and propagated for an additional microsecond. AQP1 was chosen because it presented a distinct lipid profile in its vicinity, which persisted in the atomistic simulations (Fig. S3). It is worthwhile to mention that no cholesterol flipping events occurred in the three independent 1 μs atomistic simulations with AQP1. Though a lack of flipping was consistent with reported data of a comparable membrane setup simulated with the CHARMM36 force field,^34^ it means that if one is interested in studying transleaflet cholesterol dynamics, CG simulations or atomistic force fields with different parameter sets, such as Slipids, are likely better options.^14^

In summary, our study represents an attempt to consistently evaluate and compare surface-dependant lipid profiles and cholesterol dynamics. To do so, a simple, yet physiologically representative, asymmetric biomembrane model was used, with different entities inserted. Ours findings of protein- and leaflet-dependant lipid partitioning and cholesterol flip-flop suggest that CG simulations with Martini can be effective for studying protein-lipid fingerprints.

## Acknowledgements

Work in DPTs group is supported by the Natural Sciences and Engineering Research Council (Canada). Calculations were carried out on Compute Canada facilities, funded by the Canada Foundation of Innovation and partners. DPT acknowledges further support of the Canada Research Chairs program.

