## Supplementary material for "Lipid distributions and transleaflet cholesterol migration near heterogeneous surfaces in asymmetric bilayers": SI

Table S1: System configurations.

| System | PDB | Number of lipids | | | | | | Water | Ions | Length |
|---|---|---|---|---|---|---|---|---|---|---|
| | | Upper leaflet | | | Lower leaflet | | | | | |
| | | CHOL | POPC | DPSM | CHOL | POPE | POPS | | | |
| Lipids | n/a | 78 | 57 | 57 | 78 | 84 | 43 | 2735 | 43 | 20 µs x 3 |
| Lipids*[a] | n/a | 39 | 57 | 57 | 39 | 84 | 43 | 2999 | 43 | 20 µs x 3 |
| CNT | n/a | 78 | 57 | 57 | 78 | 82 | 43 | 6305 | 43 | 20 µs x 3 |
| AQP1 | 1J4N | 78 | 57 | 57 | 78 | 82 | 43 | 6980 | 31 | 20 µs x 3 |
| KcsA | 1R3J | 78 | 57 | 57 | 78 | 96 | 48 | 6131 | 180 | 20 µs x 3 |
| OmpX | 1QJ9 | 78 | 57 | 57 | 78 | 85 | 43 | 5644 | 45 | 20 µs x 3 |
| OmpF | 3POX | 78 | 57 | 57 | 78 | 91 | 46 | 6732 | 236 | 20 µs x 3 |
| WALP23 | n/a | 78 | 57 | 57 | 78 | 84 | 43 | 3387 | 43 | 20 µs x 3 |
| AQP1 (AA)[b] | 1J4N | 78 | 57 | 57 (SSM) | 78 | 82 | 43 | 28205 | 35 | 1 µs x 3 |

[a] System containing approximately half the number of cholesterol molecules.
[b] Atomistic simulations initiated after back mapping the final configurations of the AQP1 CG systems.

Table S2: Cholesterol flip rates (flips/µs) and leaflet partitioning.

| System | Replica 1 | Replica 2 | Replica 3 | Avg ± sdev | Avg $C_{ul}/C_{ll}$ ± sdev[a] |
|---|---|---|---|---|---|
| Lipids | 0.8 ± 0.3 | 0.8 ± 0.2 | 0.8 ± 0.3 | 0.8 ± 0.02 | 1.337 ± 0.002 |
| Lipids* | 1.5 ± 0.4 | 1.4 ± 0.4 | 1.4 ± 0.4 | 1.4 ± 0.03 | 1.61 ± 0.03 |
| CNT | 0.7 ± 0.3 | 0.6 ± 0.3 | 0.7 ± 0.3 | 0.7 ± 0.02 | 1.338 ± 0.007 |
| AQP1 | 0.7 ± 0.3 | 0.7 ± 0.3 | 0.6 ± 0.3 | 0.7 ± 0.02 | 1.39 ± 0.02 |
| KcsA | 0.8 ± 0.4 | 0.8 ± 0.3 | 0.8 ± 0.3 | 0.8 ± 0.01 | 1.25 ± 0.03 |
| OmpX | 0.8 ± 0.3 | 0.8 ± 0.3 | 0.8 ± 0.3 | 0.8 ± 0.01 | 1.27 ± 0.01 |
| OmpF | 0.9 ± 0.3 | 0.9 ± 0.3 | 0.9 ± 0.3 | 0.9 ± 0.01 | 1.43 ± 0.01 |
| WALP23 | 0.8 ± 0.3 | 0.8 ± 0.3 | 0.8 ± 0.3 | 0.8 ± 0.01 | 1.32 ± 0.01 |

[a] Cholesterol ratio between the upper and lower leaflets.

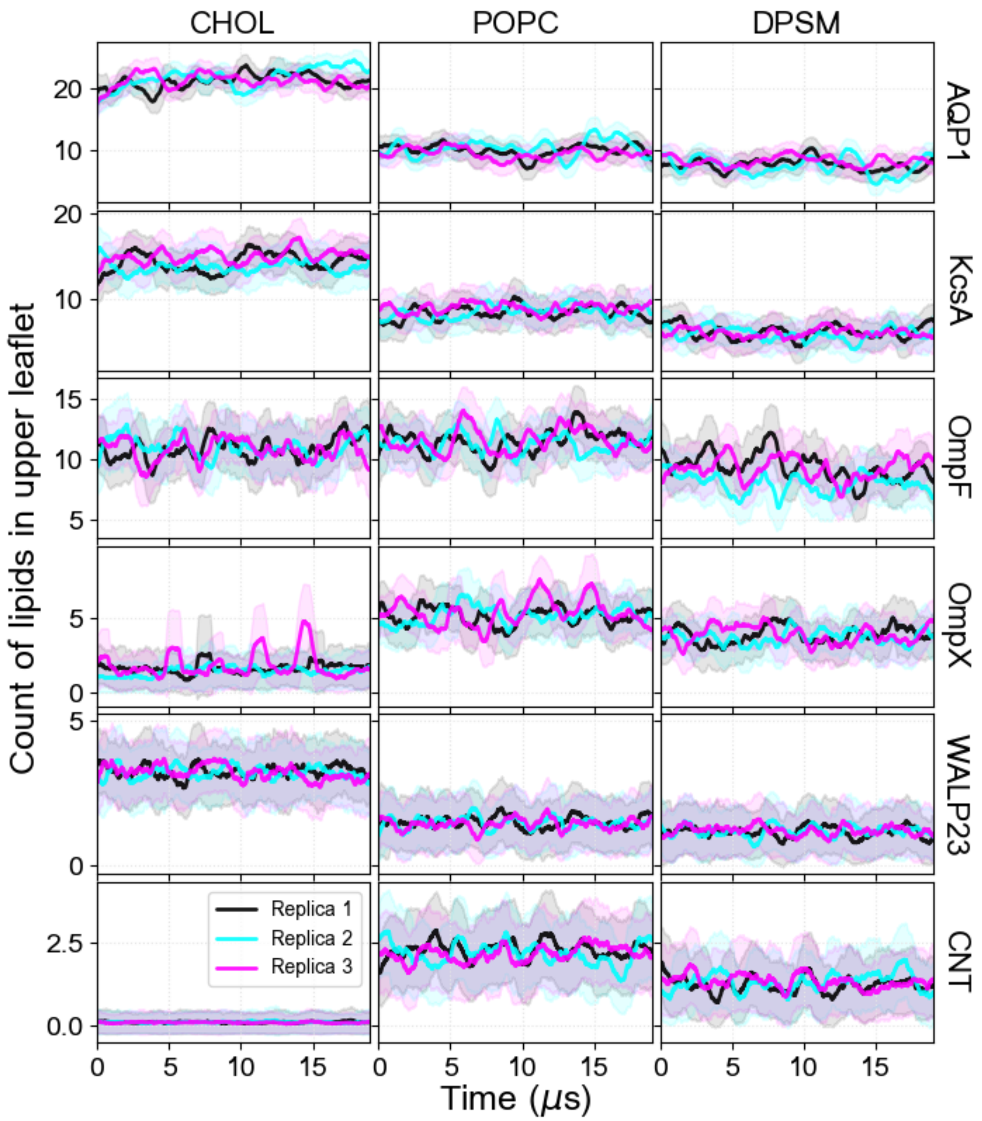


**Fig. S1 Counts of upper leaflet lipids around surfaces.** Average number of CHOL, POPC, and DPSM lipids in the upper leaflet within a 0.7 nm radius around the surface of the proteins or CNT.

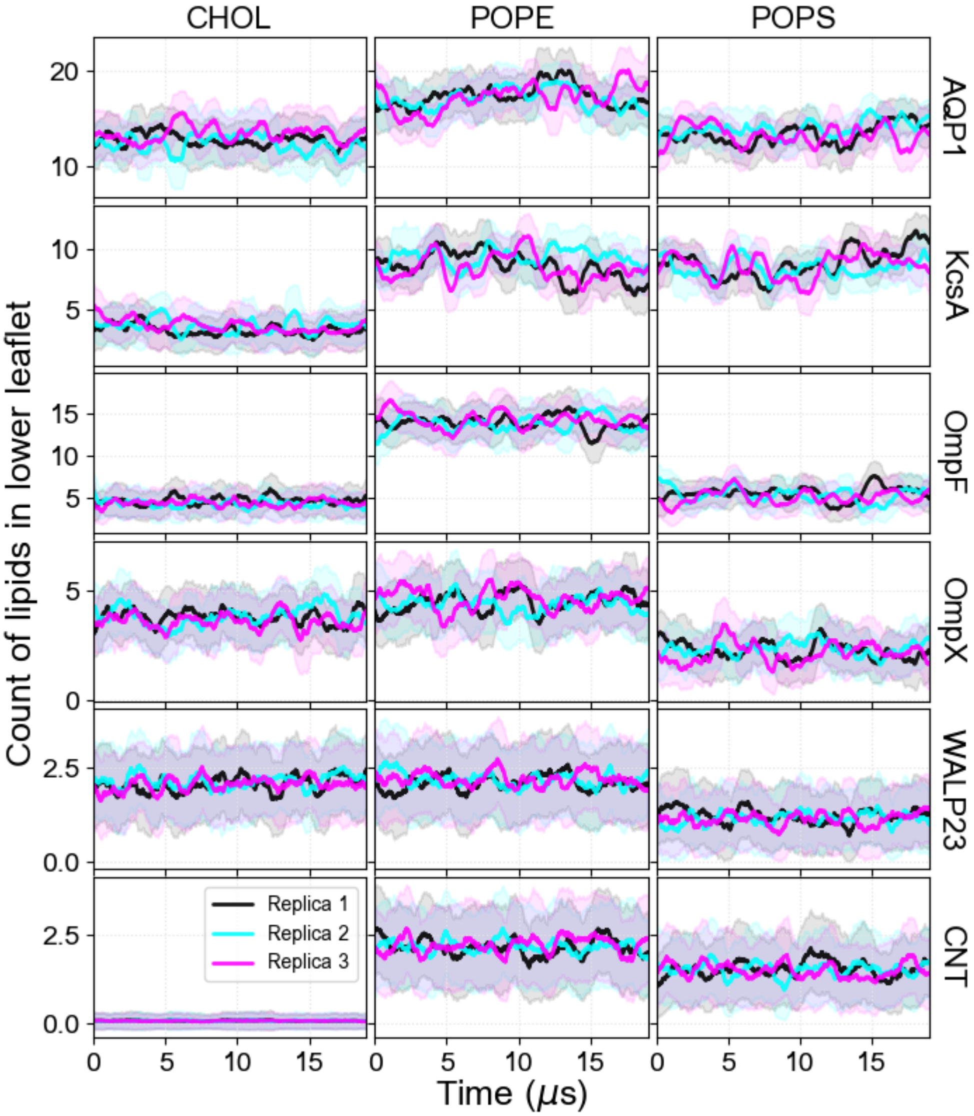


**Fig. S2 Counts of lower leaflet lipids around surfaces.** Average number of CHOL, POPC, and DPSM lipids in the lower leaflet within a 0.7 nm radius around the surface of the proteins or CNT.

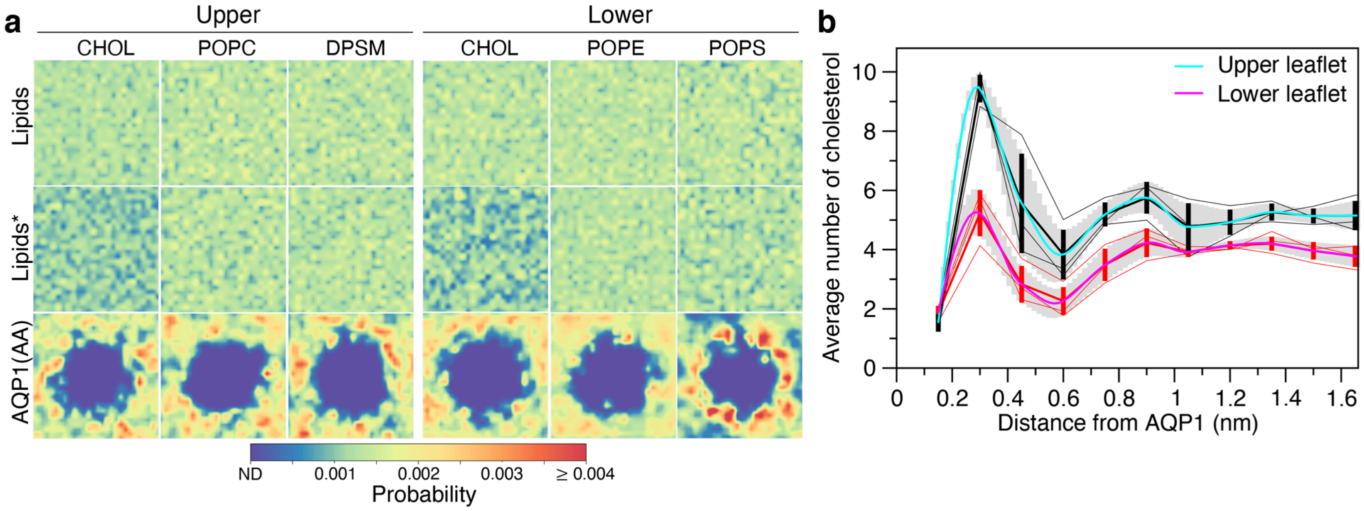


**Fig. S3 Lipid distributions. a** Probability density of CHOL, POPC, DPSM, POPE, and POPS in the upper and lower leaflets. **b** Cholesterol count as a function of distance from the AQP1 surface in the AA model. The number of cholesterol molecules were counted as a function of nearest distance from the protein to cholesterol oxygen (O3) atom for upper and lower leaflets separately. Data for individual replicas are shown in black and red for the upper and lower leaflets, respectively, with spline fits of the replica averages in cyan and magenta. Standard deviation is shown with gray error bars.

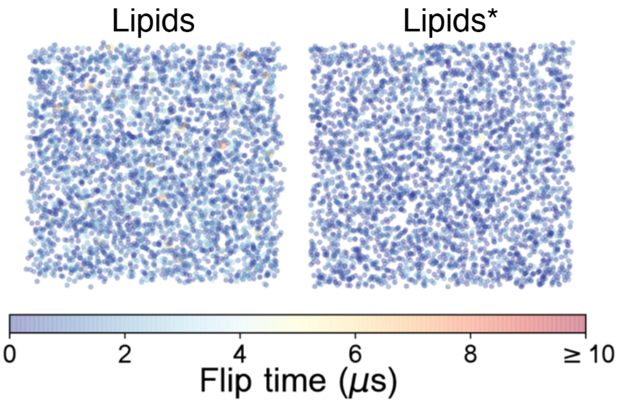


**Fig. S4 Scatter plot of cholesterol flip events and times in the lipids only systems.** Each spot represents a cholesterol flip event. Flip times are indicated by spot color and size.

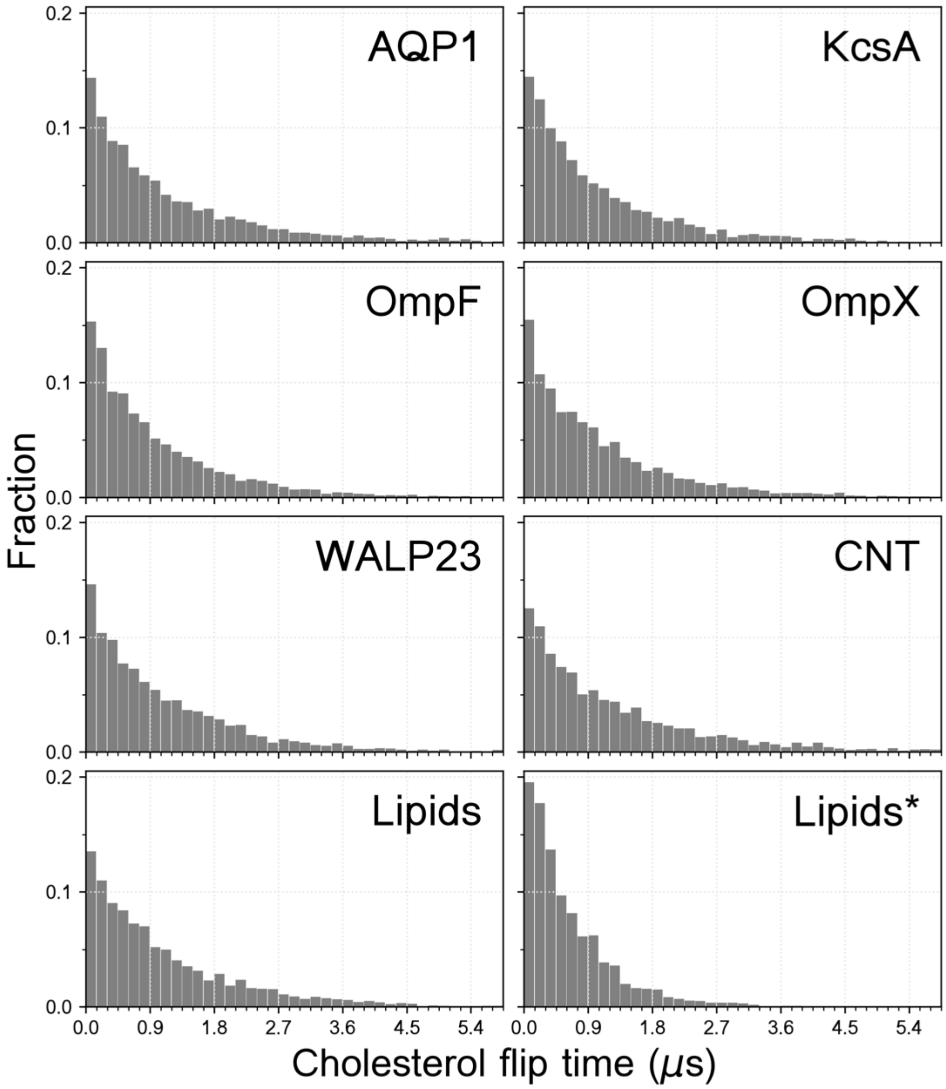


**Fig. S5 Cholesterol flip time distributions.** A bin width of 0.15 μs was used.
